# GraphMatch: Knowledge Graphs for Allogeneic Stem Cell Matching

**DOI:** 10.1101/2025.02.12.637988

**Authors:** Timothy M. Kunau

**Author notes:** Department of Computer Science and Engineering, 4-192 Keller Hall, 200 Union Street SE, Minneapolis, MN 55455.

## Abstract

Allogeneic bone marrow and umbilical cord stem cell transplants often provide the best hope for curing many patients with leukemia, lymphoma, and over 70 other diseases. Matching patients to unrelated donors requires flexible and timely searches as matching criteria change. Matching systems should scale to accommodate the diversity in patient and donor typing resolution and the growing number of donors.

We developed GraphMatch (GM), a scalable graph database solution for storing and searching variable-resolution HLA genotype markers. As a test set, we expanded the World Marrow Donor Association (WMDA) validation set based on the IPD-IMGT/HLA Database version 2.16 to create a synthetic production data set of 1 million patients and 10 million donors.

Single-patient identity search times range from 218.5 milliseconds per patient for 2 million donors to 1201.4 milliseconds per patient for 10 million donors. Search performance timing remained linear with the number of edges, even at a production scale.

In general, GM demonstrates the usefulness of graph databases as a flexible platform for scalable matching solutions.

## 1. Introduction

Allogeneic transplantation is the introduction of tissue or cells from a non-related person into the body of another [1]. Allogeneic bone marrow or umbilical cord stem cell transplants are often the best hope of a cure for many patients with leukemia, lymphoma, and over 70 other pathologies [2]. An allogeneic stem cell transplant aims to give the patient a new immune system by introducing stem cells from a donor.

Stem cell transplantation has both benefits and risks. In successful transplantation, stem cells do not attack the patient’s body but help rebuild their immune system. The new stem cells may also see cancer cells as foreign to the body and destroy them [3]. The risks include Graft versus Host Disease (GvHD) when the donor’s cells do not recognize the patient’s body as compatible and attack the host. Many factors influence the development of GvHD [4,5], including relation, gender, age, race, patient health, IL10 promoter [6], KIR genotype [7], and previous treatments, including transplants. The crucial factor in successful allogeneic transplantation remains the degree of genotype compatibility at the human leukocyte antigen locus [8].

The traditional alleles of interest are human leukocyte antigens (HLA), which are so-called because they were first discovered on white blood cells [9]. The HLA genes are the human versions of the major histocompatibility complex (MHC) [10] genes found in nearly all vertebrates. HLA antigens are essential to our immune system and defend against infectious disease, cancer, and resistance to autoimmune diseases [11]. HLA genes are critical for patient/donor matching in stem cell transplantation [12].

HLA markers are inherited similarly to other genes, half from each parent and most often as a group due to their colocation on the same chromosome [13]. Therefore, each sibling of the same parents has a 25% chance of matching the other. The MHC contains more than 220 genes, and more than 40% encode proteins involved in immune function [14]. Version IPD-IMGT/HLA 3.55.0 lists 38,909 identified alleles. [15] The human MHC is a 4Mb region on chromosome 6p21.31 [16]. As a result, allogeneic stem cell transplantation from a sibling or close relative is often the best option for treatment when a family donor is available [17].

As typing methods continue to improve and donor registries grow, we find increasing polymorphism in the MHC and persistent typing ambiguity [18]. Allele codes [19] address this imprecision by representing sets of HLA proteins most probable to code for that region. Population haplotype frequencies generated from high-resolution donor sets rely on linkage disequilibrium [20] to predict probable haplotypes. Though linkage is strongly associated with race groups [21], self-reported race designation is unreliable. Patient/donor matching tools are dynamic systems and must contend with imprecise or incomplete data. Confirmatory typing (CT) updates can improve resolution. Patients and donors are continually added and removed. However, HLA patient/donor matching methods must provide ordered matches across available race groups and variable levels of typing resolution. Traditional methods struggle to scale beyond current donor registries, which number in 10s of millions.

Accurate allele-level matching of Human leukocyte antigens (HLA) between patient and donor best predicts successful stem cell transplantation [22]. Searching the donor registries of unrelated adult stem cell donors (URD) and umbilical cord blood units (UCB) requires a programmatic method for comparing patient and donor HLA markers.

In February 2006, the National Marrow Donor Program (NMDP) released a matching algorithm called HapLogic® [23]. The program uses haplotype frequencies to predict the probability of allele-level matches. Matching improves as growing lists of donor profiles represent increasingly diverse haplotypes in the population [24]. In 2006, NMDP US adult volunteer donor profiles numbered 4.1 million [25]. Currently, NMDP supports more than 20 million donors, slightly less than half of whom are US adult volunteers [26].

HapLogic compares five markers in the HLA region of chromosome 6: HLA-A, HLA-B, HLA-C, HLA-DRB1, and HLA-DQB1 [27]. Humans are diploid, so there are two sets of five values for the patient and the same for the donor. A ten of ten (10/10) match occurs when all markers on both patient and donor profiles are equal. Lesser matches include nine of ten (9/10), eight of ten (8/10), and eight of eight (8/8) [28]. Matching requirements for URD cells are stricter than naive UCB cells [29].

We need flexible tools to capture and map the broad diversity of human histocompatibility and expand as we discover more factors that improve stem cell matching. Graph data systems can be applied to multi-omic data with structured nomenclature [30,31]. Previous work in immunology [32] has explored using the Python library NetworkX (NX) [33]. Tools such as GRIMM [34] use Neo4j [35] to represent haplotype frequencies [36,37,38] and perform haplotype-centered matching.

Graphs can represent the relationships between patients and donors, support existing data types, and expand as new criteria arise [39]. Graph databases allow adding new data types and relationships without a schema update. Graph data structures and supporting systems have emerged as powerful tools for representing relationships between numerous and diverse data sets [40]. These graphs can incorporate additional biological and demographic data to decrease match ambiguity. As we learn more about additional promoter regions and Killer-cell Immunoglobulin-like Receptors (KIR) [41], they can be incorporated into the framework, and matching accuracy will continue to improve.

This research aims to explore graph data structures to store and manipulate patient and donor allele-level genotype information so that stem cell transplant matches can be more scalable and performant than current practices allow. We developed GraphMatch (GM), a prototype graph-based system for representing patients and their associated alleles as nodes and their relationships as either “HAS_ALLELE” or “HAS_DUPLICATE_ALLELE” edges between those nodes. Donors are represented in the same way. Edges can be labeled with details illuminating the various values associated with their relationship. Basic queries for Patient/Donor matching are performed by traversing the edges from the patient, through their alleles, to the donors who share those alleles. The number (degree) of shared edges suggests appropriate donor matches. The graph representation of fundamental data supports localized searches, enabling efficient query traversals without scanning the entire dataset.

## 2. Materials and Methods

### 2.1 HLA Typing Dataset and Graph Representation

We developed an expansion of the synthetic World Marrow Donor Association (WMDA) [42,43] validation [44] set based on IPD-IMGT/HLA Database version 2.16. [45], initially including 1000 patients and 10,000 donors, with 204 distinct alleles for HLA-A, 357 for HLA-B, and 205 for DRB1.

We generated a production-sized dataset by expanding the WMDA validation set with a Python script. Each locus is represented by an array of its possible values: hla_a, hla_b, and hla_drb1, respectively. We randomly select two alleles from the array at each locus, with replacement, repeating the process across all three loci. We assembled these selections to form a synthetic typing. This process generates a random individual, which we repeat to create 1 million patients and 10 million donors.

~~~
# Select two random elements from each array
a1, a2 = random.sample(hla_a, 2)
b1, b2 = random.sample(hla_b, 2)
dr1, dr2 = random.sample(hla_drb1, 2)
# Print the pairs separated by commas
print(f”P{i},{a1},{a2},{b1},{b2},{dr1},{dr2}”)
~~~

This expanded set is created by parsing a concatenated superset of patient and donor typing from the original, extracting 766 distinct alleles into a file. This file is parsed into Python lists for each locus. Sets of synthetic patients and donors are created by selecting two values for each locus using “random.sample(hla_locus, 2)”, where each locus is in succession (‘a’, ‘b’, or ‘drb1’). The code for the expansion process is available in the GraphMatch GitHub repository https://github.com/umnkunau/GraphMatch.

### 2.2 Graph-database Design

Graph data model components include nodes, edges (relationships), and labels. Relationships between nodes can be additive and evolve. In GM, orange nodes are patients, red nodes are donors, and blue nodes are alleles. Ambiguous alleles relationships are labeled HAS_HIGHRES_ALLELE relationships. Relationships between a patient node and a representative allele node are drawn and labeled HAS_ALLELE. The patient may have a duplicate allele from each parent, and we represent that duplicate relationship as HAS_DUPLICATE_ALLELE with an additional edge. The same relationships exist for donor nodes.

In these data, alleles represent three loci in the HLA region of chromosome 6: HLA-A, HLA-B, and HLA-DRB1. Humans are diploid, so there are two sets of three values for the patient and the same for the donor.

### 2.3 Typing Ambiguity

These alleles are a collection of genes found in the human major histocompatibility complex (MHC) [46] on chromosome 6, called Human Leukocyte Antigens (HLA), and provide defense against infectious disease, cancer, and autoimmune disorders. HLA genes are critical markers for patient/donor matching in preparation for allogeneic stem cell transplantation [47] when a patient’s immune system is compromised by disease or treatment of disease. Strategies for separating and identifying HLA allele sequences in heterozygous specimens are complex and time-consuming [48]. Ambiguous haplotype assignments from serologic and sequence-based typing (SBT) [49] methods are either a process limitation [50] or a lack of phase distinction between diploid sequence reads [51].

Allele code nomenclature [52] addresses this imprecision by representing sets of HLA proteins most likely to code for that region. The NMDP allele code system manages this ambiguity by assigning a 2-5 letter code. This method was manageable when the NMDP allele code system was developed in 1990, and only a few ambiguities were thought to exist. As of April 20, 2023, 1,102,394 distinct allele codes have been assigned.

GM graphs built with the WMDA validation set contain phase and allele ambiguity. Ambiguous allele codes are expanded using the NMDP multiple-allele-code repository API [53]. The ambiguity sets are fetched from the repository for each ambiguous allele, and the relationship HAS_HIGHRES_ALLELE edges are created from the ambiguous allele node to each set member (Figure 2). GM represents ambiguous alleles with fractional edge weights, where each edge has a weight of 1 divided by the number of nodes in their high-resolution set. We know from linkage disequilibrium that not all alleles are equally likely. These weights could be informed with haplotype frequency data to fine-tune match selection despite ambiguous allele paths. Higher-resolution typing for patients and donors will help reduce typing ambiguity. As ambiguity is resolved for individual patients and donors, either through improved methods or confirmatory typing, the GM graph can be easily updated to reflect this new data.

**Figure 1.**
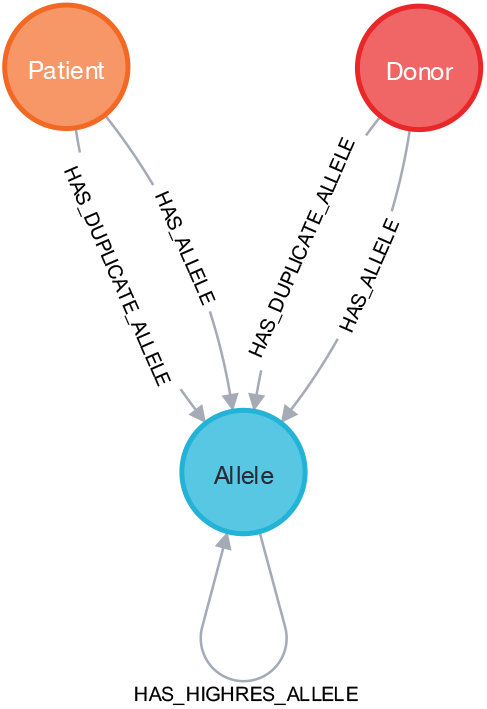
GraphMatch database schema. Patients and donors have collections of edges that describe their relationships and connect them to a set of alleles. Alleles can also indicate their type to illustrate allelic ambiguity.

**Figure 1.**
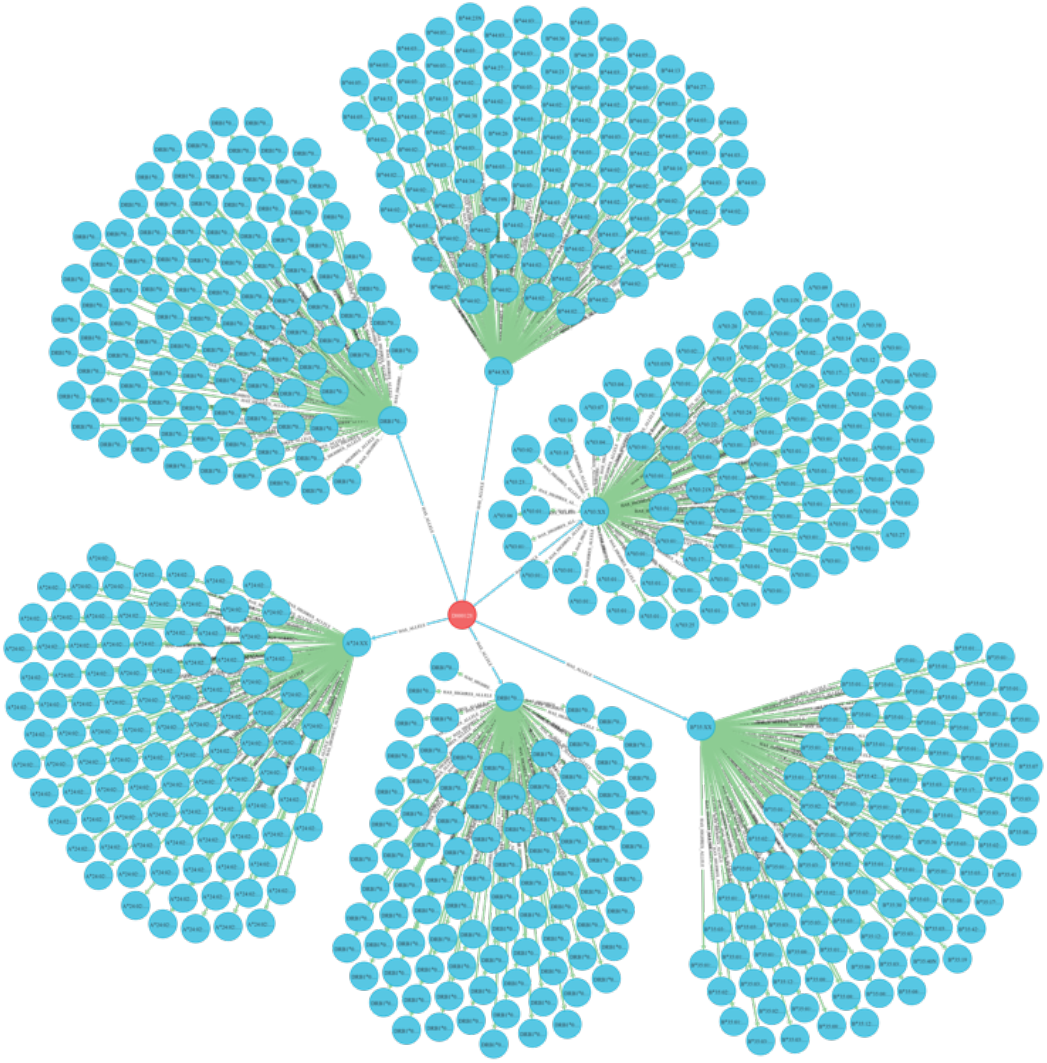
GraphMatch Donor with Allelic Ambiguity. The red circle (donor) has six relationships with blue alleles, which are ambiguously typed, each representing a set of possible high-resolution values at this locus.

### 2.4 Searching the Graph for Patient/Donor Matching

Matching tools must be deterministic, even with imprecise or incomplete data. As Sequence-based Typing (SBT) methods continue to improve and donor registries grow, we find increasing polymorphism in the Major Histocompatibility Complex (MHC) and persistent typing ambiguity.

Patient/Donor matching begins with a unique patient node and is performed by traversing the edges from that patient to available donors through their shared alleles. A basic Cypher query for patient ‘P000128’, shown below, locates all the attached Alleles with either label (HAS_ALLELE or HAS_DUPLICATE_ALLELE), and uses those alleles to find donors with more than 3 shared alleles. The query returns a prospective donor list of shared relationships ordered by degree. The higher the degree, the stronger the match. GM orders that list of donors and limits it to 10. This is the GraphMatch Cypher query for an identity match:

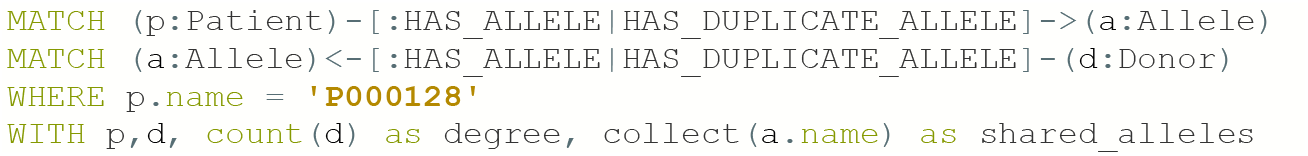
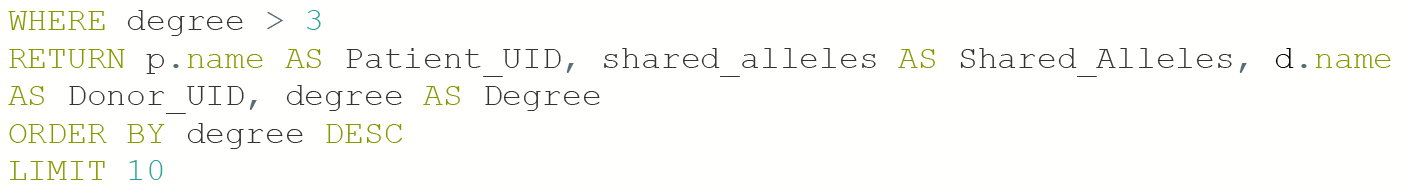

The Neo4j console displays a table with results in the same order as the query RETURN statement.

A six-of-six (6/6) match occurs when patient and donor profiles share all six of the six compared alleles. This count is expressed by the degree of shared edges. Matches may include five of six (5/6), four of six (4/6), and three of six (3/6). Mismatches are discovered by collecting the allele sets of shared donors and subtracting the alleles shared with the patient.

Matching is performed on the local network of connected neighbors and does not require traversal of the entire graph. The resulting graph may look like the subset of nodes and edges in Figure 3.

**Figure 3.**
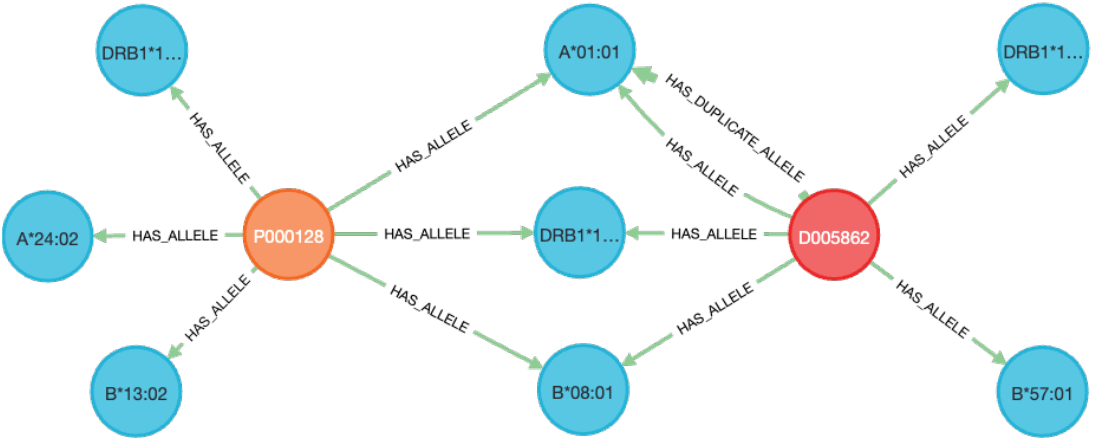
Patient and Donor match through three shared alleles. P000128 shares three alleles with D0005662. The donor in this match has a duplicate A*01:01 allele. Different scoring methods may count these duplicate allele matches as better or worse.

The query example is a search for identity matching and relies heavily on reliable typing for patients and possible donors. When typing is less reliable and ambiguity is introduced, we can modify the query to allow paths through the ambiguity sets in the Allele graph. Alternatively, we can use haplotype frequencies to identify probable values for each ambiguous locus and rerun the search. These choices could be selected in the clinical-facing tool and tagged to indicate match imprecision.

Matching requirements for unrelated donor (URD) cells are stricter than for naive unrelated cord blood (UCB) cells [54]. The WMDA validation set does not state the origin of the donor cells. However, a donor repository would have this information, and it could easily be added as a property of the donor node.

### 2.5 GraphMatch Output and Benchmarking

We developed GM within the Neo4j console, as Table 1 displays. This allowed rapid prototyping of the query and visualization of graph components (Figure 2) and results (Figure 3). Ultimately, GM could be a component of a transplant management system. This would require network API access and a set of parsable results. We developed this network interface to benchmark our GM instance with our expanded WMDA data set.

**Table 1.** GraphMatch top 10 identity match results from the Neo4j console.

| Patient_UID | Shared_Alleles | Donor_UID | Degree |
| --- | --- | --- | --- |
| "P000128" | ["DRB1*15:01", "DRB1*13:02", "B*08:01", "A*01:01", "A*01:01"] | "D2727485" | 5 |
| "P000128" | ["DRB1*15:01", "DRB1*13:02", "B*08:01", "B*13:02", "A*01:01"] | "D6828082" | 5 |
| "P000128" | ["DRB1*15:01", "B*08:01", "B*08:01", "A*01:01", "A*24:02"] | "D4098369" | 5 |
| "P000128" | ["DRB1*15:01", "B*08:01", "B*08:01", "A*01:01", "A*01:01"] | "D007196" | 5 |
| "P000128" | ["DRB1*15:01", "DRB1*15:01", "B*08:01", "A*01:01", "A*24:02"] | "D4319338" | 5 |
| "P000128" | ["DRB1*15:01", "B*08:01", "A*01:01", "A*01:01"] | "D9842357" | 4 |
| "P000128" | ["DRB1*15:01", "B*08:01", "A*01:01", "A*24:02"] | "D9875791" | 4 |
| "P000128" | ["DRB1*15:01", "DRB1*13:02", "B*13:02", "A*24:02"] | "D9950037" | 4 |
| "P000128" | ["DRB1*15:01", "DRB1*15:01", "A*01:01", "A*01:01"] | "D9715373" | 4 |
| "P000128" | ["DRB1*15:01", "B*08:01", "B*13:02", "A*01:01"] | "D9894728" | 4 |

Benchmarking was performed through a locally run Python script (graphmatch.py) where all network traffic is through the loopback interface to minimize the effects of network variability. The Python script uses the GraphDatabase class of the Neo4j module and ingests connection details from a local .env file. With post-processing, non-ambiguous match results can be reported on a single line (Figure 4) in the following order: **degree** of matched alleles, **unmatched patient alleles, patient identifier**, the **alleles shared** between patient and donor, **donor identifier**, and **unmatched donor alleles**.

**Figure 4.**
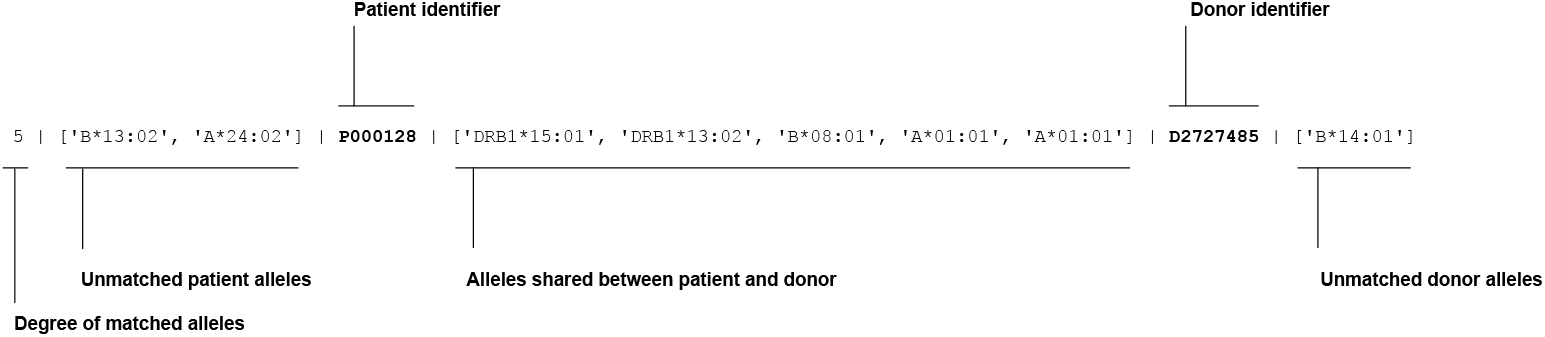
GraphMatch detail result from Python. This single line of output represents the match criteria for P000128 with D2727485. Processing after the match is found allows our benchmarking script to collect and display unmatched alleles for both the patient and donor.

The search set was supplied as a file on the command line. Sets of Patient identifiers were offered for serial searches in sets of 1, 10, 100, and 1000. Parallel searches were performed in 10 groups of 100 Patients each. Timing results were gathered from the UNIX “time” command output and represent the “real” time result, not the “user” or “sys” time. For this benchmark, the number of searches performed for each patient may vary from 1 to 12, according to the following procedure. The first search finds all the connected donors with a degree greater than 3, orders the list by descending degree, and limits the findings to the top 10. If zero connected donors are found, no other search is performed. If between 1 and 10 donors are found, an additional search is performed to discover all the alleles for that patient and each donor. The code for this command-line search is available in the GraphMatch GitHub repository https://github.com/umnkunau/GraphMatch.

## 3. Results and Discussion

We developed GraphMatch (GM), which stores and facilitates patient/donor matching for stem cell transplantation based on shared alleles. GraphMatch leverages a graph database platform, Neo4j, instead of a traditional Relational Database Management System architecture. RDMSs rely on normalized tables where relationships emerge through complex joins. While effective for specific scenarios, this approach can suffer from performance bottlenecks when dealing with network structures like patient/donor relationships.

### 3.1 Graph Databases as a Solution for Patient/Donor Matching

Graph databases offer a compelling alternative to RDBMS’ by modeling relationships as fundamental data objects. While relational systems are composed of tables where data are arranged in rows and columns, graph databases are a collection of nodes (vertexes) with edges (relationships) connecting one node to another.

Traditional relational systems use joins to find connections between data in tables. However, these joins can be slow when dealing with large datasets. In contrast, GraphMatch graph queries only navigate through relevant parts of the network, providing consistent performance even with substantial amounts of data. This method avoids combinatorial explosion [55], a common performance issue in relational systems. The fundamental mapping of connections between patients and donors is central to patient-donor matching, allowing for efficient traversal of relationships. This mapping enables real-time retrieval of relevant connections without searching an entire database (see Methods).

In GraphMatch, donors and patients are stored as nodes in the graph, with edges connecting these patient and donor nodes to shared alleles relevant for transplant matching. Given this graph, queries to the system do not rely on traditional indexing for patient/donor matches. Instead, a global index is used to find the initial patient node; all queries are local to the patient node. This condition is called “index-free adjacency” (IFA) [56]. GM searching is limited to the edges between the selected patient, their alleles, and the connected donors (see Methods).

Production patient/donor matching systems actively employ interactive queries and batch operations for efficient patient/donor matching and dataset management. Interactive matching functions are primarily utilized during clinic hours with a human in the loop, while batch functions perform coordinated updates and expansions in a scheduled manner. On GM, batch updates can seamlessly coexist with interactive matching processes.

### 3.2 Benchmarking Design

To test the utility of the GraphMatch system in supporting patient/donor matching on a production-scale donor set, we established a synthetic cohort with genotypes. We carried out a series of benchmarking experiments. Specifically, we used the WMDA validation data (see Methods) to derive a synthetic set of 1 million patients and 10 million donors and their associated HLA alleles. We designed benchmark experiments exploring various search configurations, which were then executed on two Neo4j hosts (Table 2).

**Table 2.** GraphMatch tuning sets, conditions, and systems. System 1 is a 4-CPU desktop system with 32GB of RAM and an internal SSD. System 2 is a 24-CPU server-class system with 128 GB of RAM and a JBOD chassis.

| Counts and Conditions | System 1<br>(4 CPU/32 GB RAM) | System 2<br>(24 CPU/128 GB RAM) | Comment |
| --- | --- | --- | --- |
| 1 search | 2.5s | 4.9s | 1 group of 1 |
| 10 serial searches | 26.5s | 49.9s | 1 group of 10 |
| 100 serial searches | 195.9s | 372.8s | 1 group of 100 |
| 1000 serial searches | 2152.4s | 3814.5s | 1 group of 1000 |
| 10 parallel threads of 100 | 1201.4s | 1038.9s | 10 groups of 100 |

Performance tuning is a multi-variate problem. In our example, System 1 (a compact desktop system) and System 2 (an older server-class system) have broad architectural differences. System 1 has limited CPU and RAM resources to manage operating systems, applications, and interactive user services. System 2 is designed to support interactive network services and operate under heavy load, with slower CPUs and larger RAM resources.

We began with 1, 10, 100, and 1000 serial searches across the same collection of a 10 million donor set. Parallel searches of 10 groups of 100 were also tested.

### 3.3 Results of Benchmarking

To illustrate GM’s scaling performance, we ran 20 sets of 1000 serial searches on System 1, with 1 million patients, as we scaled the donor set from 2 to 10 million donors (Figure 5). This iterative approach allowed us to evaluate the system’s scalability and efficiency under production-scale conditions. These searches display a linear scaling in single-threaded search time from 2 to 10 million donors. Single-patient search times ranged from 218.5 msec/patient for 2 million donors to 1201.4 msec/patient for 10 million donors. These times represent practical production searching at scale. Performance timing is linear with the number of edges (Figure 6).

**Figure 5.**
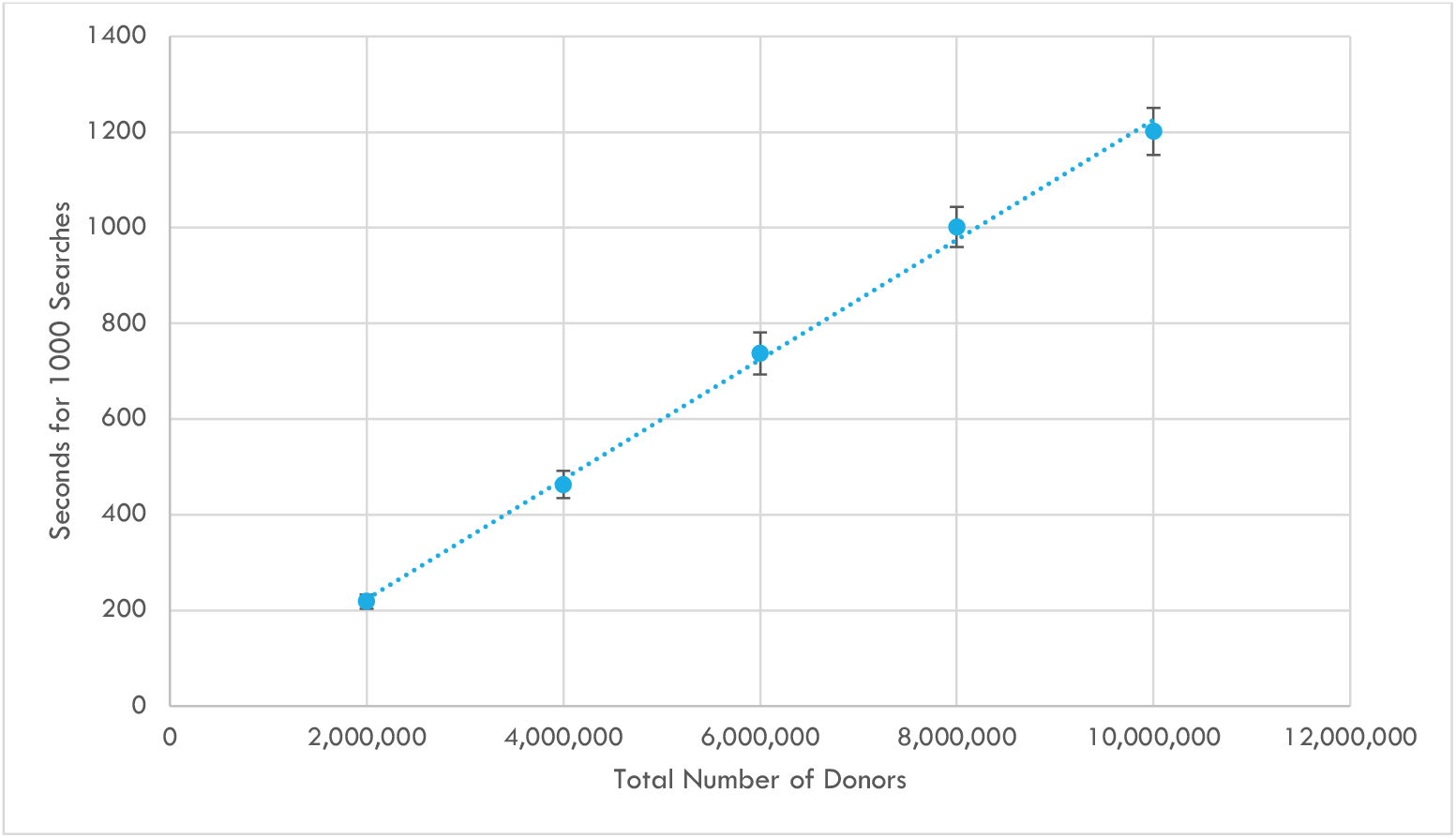
Search runtime as a function of Donors. Each data point represents the average of 20 sets of 1,000 GraphMatch searches as donors increase from 2 million to 10 million. The error bars at each data point represent the standard deviation.

**Figure 6.**
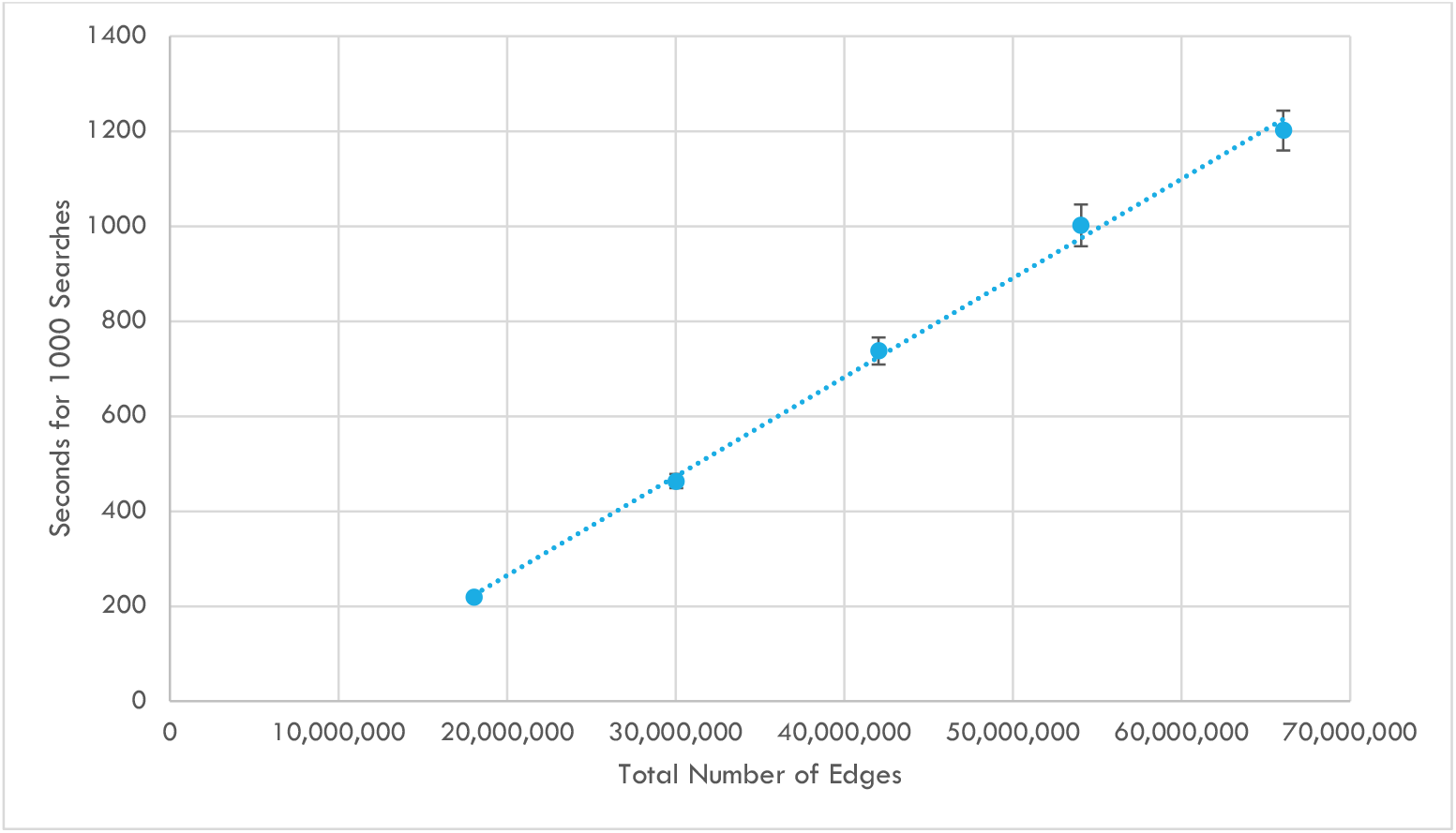
Search runtime as a function of Edges. Each data point represents the average of 20 sets of 1,000 GraphMatch searches as edges from 18,032,631 to 66,032,637. The error bars at each data point represent the standard deviation.

The results presented in Figures 5 and 6 support these observations. As the total number of donors and edges in the database increases, the average time required to perform 1,000 GraphMatch searches remains remarkably stable, demonstrating the scalability and efficiency of the underlying graph data structure.

The query benchmarks in Table 2 illustrate computational speed differences as search sets grow. System 1 is consistently faster in single-threaded patient/donor searches until those searches are distributed across multiple processors. In this parallel search example, ten parallel searches are offered to System 1 and its 4 CPUs, while ten parallel threads are offered to System 2 and its 24 CPUs. In the case of the former, there are more threads than CPUs, and performance suffers when threads wait for processing. In the latter, there are fewer threads than CPUs, and even though these CPUs are slower, the threads do not wait for processing. System 2 is faster under 10 groups of 100 parallel searches, though its CPUs are slower computationally. System 1 would benefit from parallel processing loads; in serial tests, the faster CPU in System 1 was consistently more performant. For this reason, we chose 1000 serial searches as the test set and System 1 as the platform for serial benchmarking to mitigate this differential.

### 3.4 Scalability

Production graph database platforms, including Neo4, are scalable, supporting billions of nodes and relationships. Neo4j can scale both vertically and horizontally: vertically by building a more extensive single system with multiple CPUs and server-class memory and horizontally by creating clusters of similarly sized systems and partitioning (sharding) both the data and GM processing across the topology. Software performance tuning is a balancing act to support the system’s competing needs.

A production environment will have a mixture of maintenance updates and patient/donor matching queries. In some organizations using single-system search platforms, maintenance updates must occur ‘after business hours.’ Service expectations increasingly demand 24-hour availability. To balance these varied loads, a collection of modestly resourced systems will outperform a single monolithic system and continue to scale beyond the limits of a monolithic system. GM is built on a graph-based platform to support interactive patient searches in web time, even as the donor domain grows.

Software may also have hard-coded limitations. The Neo4j Graph Data Science Library - Community Edition (GDS CE) software, used in this GM example, limits concurrency to 4. In contrast, the Neo4j Graph Data Science Library - Enterprise Edition (GDS EE) is unlimited.

### 3.5 Platform Flexibility

GraphMatch demonstrates a flexible platform that can capture and map the broad diversity of human histocompatibility and expand as we discover more factors that improve stem cell matching. Our method offers a more cost-effective solution by avoiding the need for massive computing resources. Our graph-based model allows for efficient memory usage, traversing adjacent alleles and associated donors, avoiding the need for unbounded table joins. This efficiency contributes to the platform’s overall performance and responsiveness. The production timings were performed with the Community Version of Neo4j on a modest desktop system (Table 2). These capabilities reduce the necessity for larger single-system platforms and provide a practical alternative for organizations with limited resources.

Identifying and modeling an entire domain when assembling a data collection is difficult. The schema will likely expand as data becomes available and the domain becomes increasingly understood. In the case of patient/donor stem cell matching, these changes demand a flexible and expressive model to represent the relationships between patients, donors, and match criteria. GraphMatch allows you to begin with a minimum viable product and grow your system over time. Our platform supports incremental modeling and can expand as you become aware of additional relationships and your match requirements change.

GraphMatch’s adaptabilities extend to the ease of integrating various data sources and types. We adopted the industry standard Allele naming ontology [57]. Given the patients’ and donors’ associated alleles, we generated shared relationships for stem cell matching. Our approach supports optional matches for ambiguous data to address a common challenge in patient-donor matching. This feature allows cascading searches, accommodating cases where more precise matches are unavailable as richer data becomes available. The alleles in the GraphMatch schema may be augmented with additional relationships supporting a broader search domain.

GM competes favorably by offering comparable functionality and without proprietary black-box methods. GM can mimic the functionality of other methods, such as haplotype-based matching [58], demonstrating its versatility and underscoring its capacity to encompass various search methodologies within a unified framework. This flexibility supports the emulation of various search methods, allowing for the parallel display of match results using different methods from the same underlying data.

### 3.6 Limitations

Graph databases are not appropriate for all queries. If a query demands entities in the model, such as binary large objects (BLOBs), character large objects (CLOBs), or large text fields, Relational Database Management Systems (RDBMS) are better designed to handle this data. It is common practice to build graph-based systems with supporting RDBMS systems to manage supporting data. Similarly, match scores are often calculated once the prospective donor set is collected. Scores may affect the way results are displayed graphically for the user. These scores and displays may occur external to the graph database.

Combinatorial expansion of the WMDA dataset will create more duplicates and represent less diverse sets than in nature. However, production donor bases do not necessarily represent nature. Donor drives at universities will produce donors between 18 and 25. Donor drives in different regions of a country may produce clusters of similar haplotypes. [59] Donors are collections of communities where and when they are gathered. Our expanded dataset is likely unrealistic and results in more over-matched donors than might appear in organic donor populations of a similar size. Additionally, it likely represents a more challenging test than an organic production donor set, placing additional load on the SORT and LIMIT steps of the query.

### 3.7 Future Research

There are a number of directions for future extensions of this work. For example, GM could support other Genotype representations. Genotype List (GL) Strings [60] would be a valuable improvement. GL Strings express loci and alleles, phases, haplotypes, and possible genotypes. The imputation of higher-resolution patient and donor typing [61] is based on publicly available haplotype frequencies. Edges could be added to the GM schema to represent these relationships and support novel search methods.

GM could incorporate haplotype imputation. Population haplotype frequencies generated from high-resolution donor sets rely on linkage disequilibrium [62] to predict probable haplotypes.

These frequencies could be used as edge weights to sort allele-level matches. Traditionally, matching considers self-reported race groups [63], even though self-reported race designation is unreliable.

Ultimately, GraphMatch provides a practical alternative for organizations with limited resources by offering a flexible, scalable, and performant patient/donor matching system that can grow over time to accommodate new findings and requirements in human histocompatibility.

## 4. Declaration of Competing Interest

The author declares that they have no known competing financial interests or personal relationships that could have appeared to influence the work reported in this paper.

## 5. Acknowledgments

The University of Minnesota, Minneapolis, MN, provided data systems and support.

## Abbreviations

GM: GraphMatch
GvHD: Graft versus Host Disease
HLA: human leukocyte antigen
MHC: Major Histocompatibility Complex

## Notes

https://github.com/umnkunau/GraphMatch

### Competing Interest Statement

The authors have declared no competing interest.

